# Regional Brain Entropy During Movie-watching

**DOI:** 10.1101/2024.06.12.598767

**Authors:** Dong-Hui Song, Ze Wang

## Abstract

Brain entropy (BEN) measures the irregularity or unpredictability of brain activity. Our previous studies have showcased the neurocorrelates and disease sensitivity of regional BEN. Task performance and neuromodulations as well as medications can modulate regional BEN. Movie-watching engages multiple-domain cognitions and complex brain activities mimicking the complex real world and has been increasingly used to probe brain state dynamics. The sustained multiple-domain cognitive activities would cause sustained changes to regional BEN, which however remains unknown. The purpose of this study was to address this gap. We first aimed to examine whether the fMRI-derived regional BEN represents a stable brain activity index during repeated measurement with or without movie-watching. We then assessed the movie-watching-induced regional BEN changes compared to resting state and regional BEN differences between different movie clips. we found higher reliability during movie-watching and movie-watching induced lower BEN in the sensory cortex and higher BEN in the association cortex compared to the resting state. We replicated and validated all findings in test-retest clips. This work systematically evaluated the regional BEN during movie-watching, providing a new direction for exploring brain activity under natural paradigms using BEN.

## Introduction

Entropy is a concept derived from thermodynamics, used to characterize the degree of disorder [1]. According to the second law of thermodynamics, entropy increases over time in a closed system and cannot decrease, eventually reaching a state of chaos. Subsequently, Shannon used entropy to quantify uncertainty and information content in information theory [2]. The human brain, considered one of the most complex systems in the world, is believed to possess a high degree of self-organization [3, 4], an organization in which initial disorderly systems interact locally between various components, forming some form of overall order and this process of organization can spontaneously occur without the need for external control when energy supply is adequate. The human brain continuously interacts with the external environment, exchanging information while consuming a significant amount of energy to thrives to maintain its entropy balance to keep its self-organization and to achieve functional flexibilities as a complexity dynamic system [5-8].

Brain entropy (BEN) can be assessed using various signals of brain activity, including EEG [9, 10], MEG [11], and fMRI [12], employing different algorithms for quantification such as Shannon entropy [2, 13, 14], approximate entropy [15], and sample entropy [12, 16]. Due to its higher spatial resolution, fMRI has gained increasing attention in recent years for BEN mapping [12]. Additionally, sample entropy can characterize the entropy of shorter physiological time series [16, 17], further enhancing the advantages of using fMRI to assess BEN. Our previous studies have showcased the neurocorrelates in large cohorts and the disease sensitivity of regional BEN [18-21]. Similarly, regional BEN can also be modulated by non-invasive brain stimulation, medication, and non-pharmacological therapies, reflecting its sensitivity to neuroplasticity [19, 22-25]. More importantly, recent studies have shown lower resting brain entropy is associated with stronger task activation and deactivation, reflecting the close relationship between regional BEN and cognitive brain activity [26].

Naturalistic stimuli, such as movie-watching, offer a greater constraint compared to resting-state and provide more ecological relative to than traditional task designs [27]. Movie-watching engages multiple-domain cognitions and complex brain activities mimicking the complex real world and has been increasingly used to probe brain state dynamics [28-31]. The sustained multiple-domain cognitive activities would cause sustained changes to regional BEN, which however remains unknown. The purpose of this study was to address this gap. We first aimed to examine whether the fMRI-derived regional BEN represents a stable brain activity index during repeated measurement with or without movie-watching. We then assessed the movie-watching induced regional BEN changes.

## Results

### The better test-retest reliability in regional BEN during movie-watching compared to the resting state

We employed intra-class correlation coefficients (ICC) to assess the reliability of regional BEN between resting-state and movie-watching states. We separately evaluated the reliability of scans on the first (day 1) and second days (day 2) and similarly assessed the reliability between the four scanning runs. The results shown that both resting-state and movie-watching BEN exhibited mild to high reproducibility (r > 0.3) across the whole brain. However, compared to the resting state, BEN during movie-watching demonstrated relatively higher ICC in the ventromedial prefrontal cortex (VMPFC) and posterior cingulate cortex (PCC) (p<0.05), indicating greater reproducibility in these regions during movie-watching (Fig 1).

**Fig 1.**
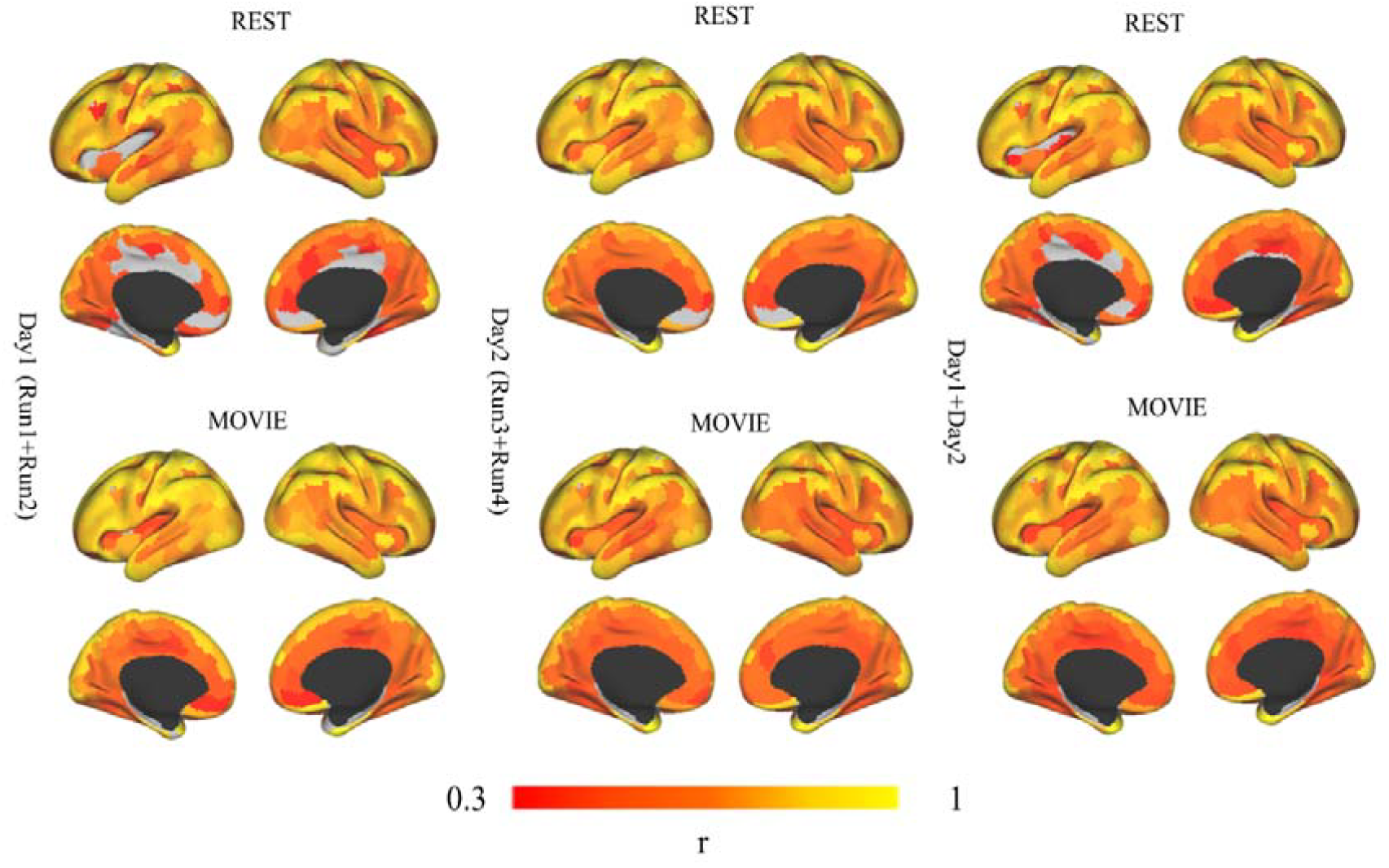
ICC of BEN for resting-state and movie-watching. Left: The upper row represents the ICC of resting-state BEN, and the bottom row represents the ICC of movie-watching BEN on day 1. Middle: The upper row represents the ICC of resting-state BEN, and the bottom row represents the ICC of movie-watching BEN on day 2. Right: The upper row represents the ICC of resting-state BEN, and the bottom row represents the ICC of movie-watching BEN for all four runs. Colorbar indicates the correlation coefficient r value.

### Movie-watching reduced regional BEN in the sensory cortex but increased entropy in the association cortex

Compared to the resting state, movie-watching showed lower BEN in the sensory cortex including the visual cortex and temporal cortex, but higher BEN in the association cortex including default mode network (DMN) (PCC/precuneus (PCC/PCu), angular gyrus (AG), dorsomedial prefrontal cortex (DMPFC)) and frontoparietal network (FPN) (dorsal-lateral prefrontal cortex (DLPFC) and posterior parietal cortex (PPC)) (Fig 2a). Functional decoding using Neurosynth found that this difference is mainly positively related to memory processing, such as “retrieval”, and “memory” and negatively related to audio-visual functions, such as “visual”, and “auditory” (Fig 2a).

**Fig 2.**
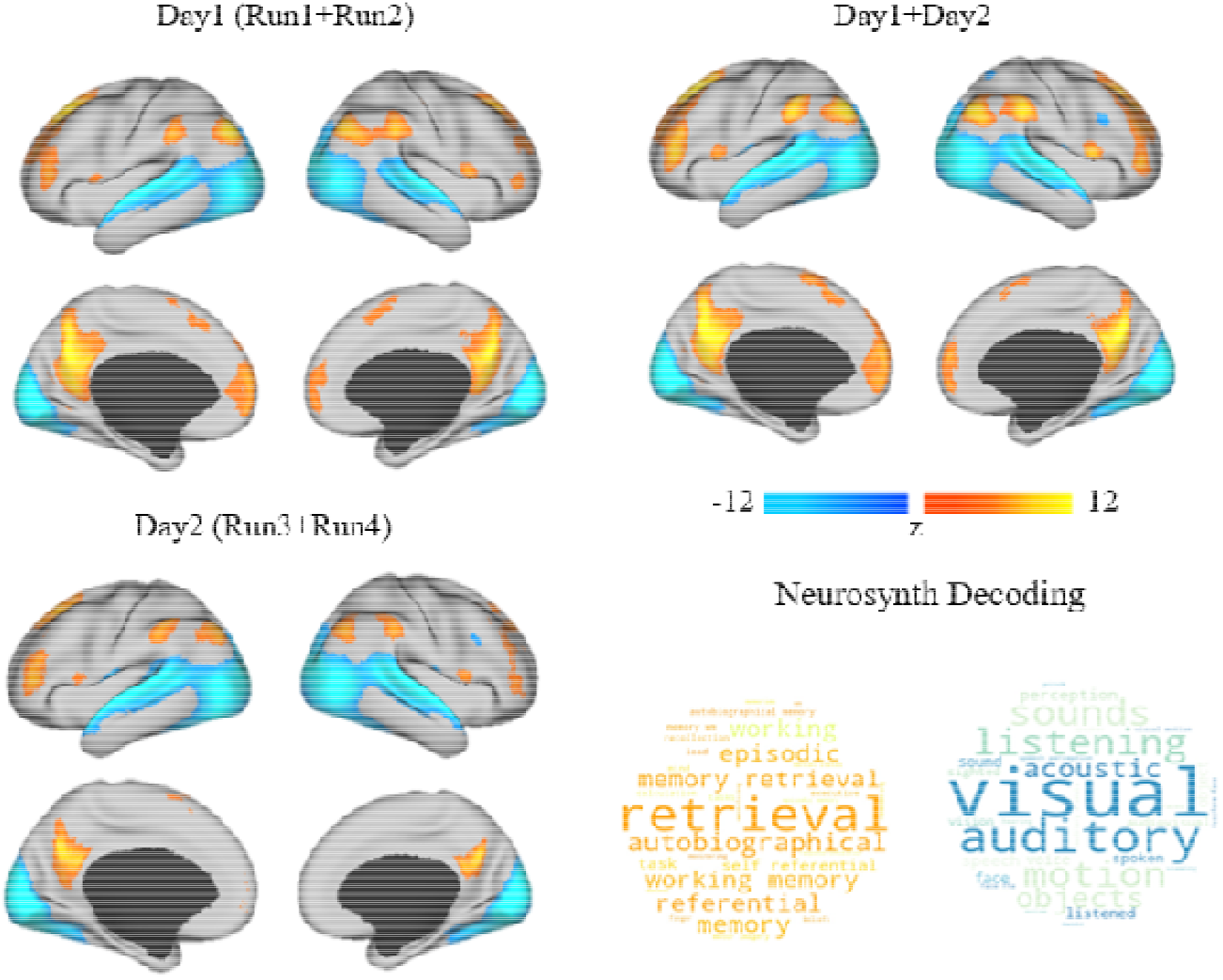
Differences of BEN between movie-watching and resting-state (FWE _corrected_ p<0.05). Left: The upper row represents differences of BEN between movie-watching and resting-state in day1, and the bottom row represents differences of BEN between movie-watching and resting-state in day2. Right: The upper row represents differences of BEN between movie-watching and resting-state for all runs, and the bottom row represents Neurosynth decoding from unthresholded statistical maps of differences of BEN for all runs, hot color means positive correlation, blue color means negative correlation, font size indicates correlation strength. Colorbar indicates z value, hot color means higher BEN in movie-watching, cool color means higher BEN in resting-sate.

### Differences of movie clips were captured by regional BEN

To confirm whether regional BEN can capture differences between movie clips, we assessed differences between three clips within each run. Additionally, to mitigate potential time effects that could influence BEN differences across scans acquired at different times, we also evaluated differences between corresponding TRs within each run during resting-state. The results indicated that clips between each run yielded unique BEN differences (Fig 3a), while in the resting-state, consistent differences were observed across runs in the visual and motor cortices (Fig 3b).

**Fig 3.**
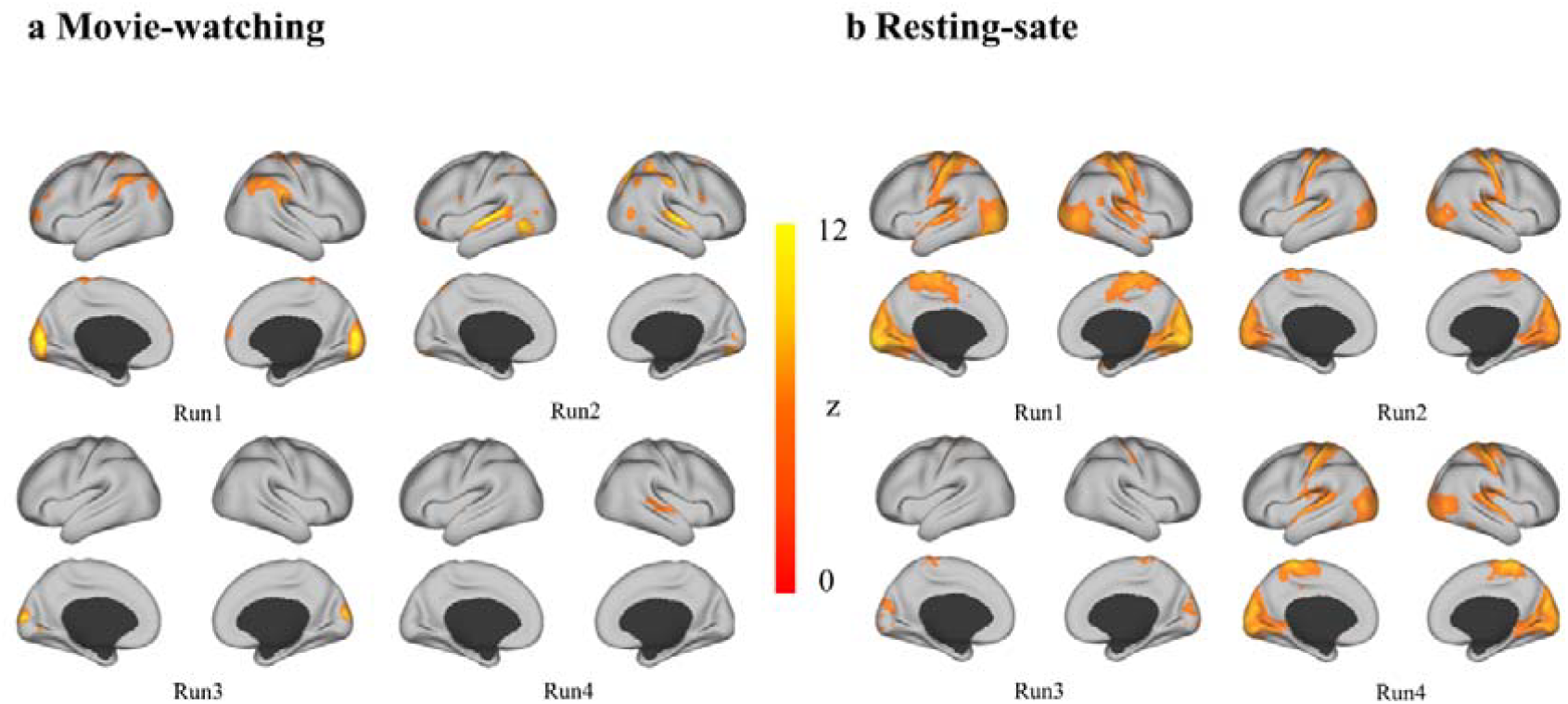
Differences of clips were captured by regional BEN (FWE _corrected_ p<0.05). a) The differences in BEN among the three clips during movie-watching in four runs. Following the clockwise direction, they are in run 1, run 2, run 3, and run 4, respectively. b) The differences in BEN for corresponding TRs during the resting state in run1, run2, run3, and run4. Colorbar indicates the z value, hot color means the degree of difference.

### Validation and robustness analyses

Using the test-retest clips, we further confirmed that BEN exhibits higher test-retest reliability during movie-watching compared to the resting state (Fig 4a). Moreover, movie-watching induced higher BEN in the association cortex and lower BEN in the sensory cortex (Fig 4b). Similarly, in the test-retest clips, we did not observe significant differences in BEN between different runs, further indicating that BEN differences in movie clips does not arise from noise.

**Fig 4.**
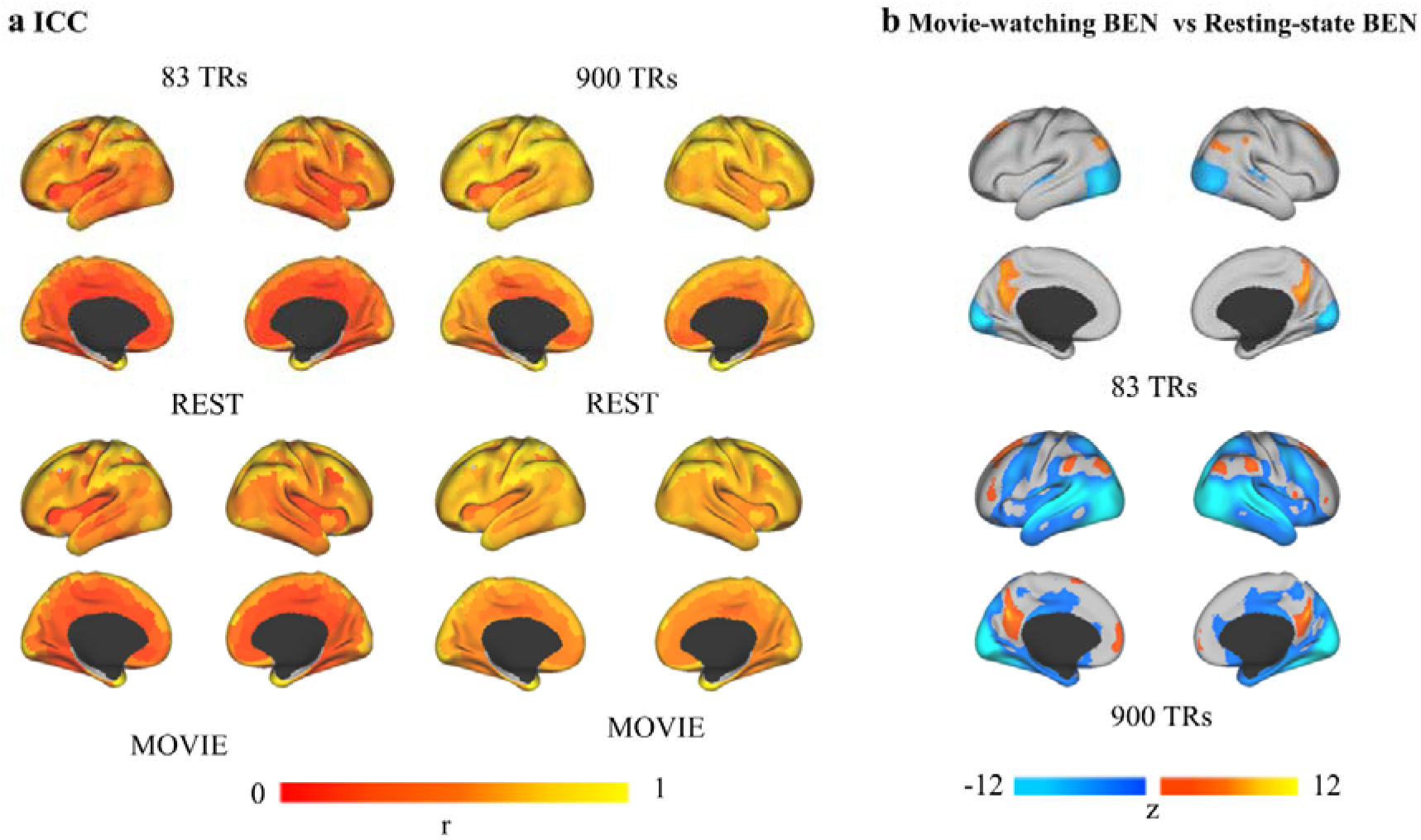
Validation and robustness analyses for reliability and movie-watching induced BEN. a) Left: The upper row represents the ICC of resting-state BEN, and the bottom row represents the ICC of movie-watching BEN during test-retest clips from four runs. Right: The upper row represents the ICC of resting-state BEN, and the bottom row represents the ICC of movie-watching BEN during 900 TRs. Colorbar indicates the correlation coefficient r value. b) The upper row represents differences in BEN between movie-watching and resting-state for all runs during test-retest clips (FWE _corrected_ p<0.05). The button row represents differences in BEN between movie-watching and resting-state for all runs during 900 TRs (FDR _corrected_ p<0.05). Colorbar indicates z value, hot color means higher BEN in movie-watching, cool color means higher BEN in resting-sate.

## Discussion

We found higher reliability in regional BEN during movie-watching compared to the resting state. This finding is consistent with previous results based on other measurement metrics [27], such as Wang et al found that the reliability of connectivity and graph theoretical measures of brain networks is significantly improved during movie-watching over resting-state [32] and ICC evaluation based on functional connectivity (FC) revealed that the ICC of movie-watching FC across different movies was comparable to that of resting-state FC across repeated scans using HCP dataset [33]. Specifically, higher test-retest reliability in BEN during movie-watching is primarily concentrated in VMPFC and PCC, core regions of DMN. The DMN primarily supports complex internal mental state, and the higher test-retest reliability in these regions may be due to the relatively consistent internal brain activity during movie-watching. In contrast, the less constrained nature of resting-state, where internal mental state fluctuations are more random, leading to relatively lower reliability.

we found lower BEN in the sensory cortex and higher BEN in the association cortex during movie-watching compared to the resting state. Lower BEN in the sensory cortex can be attributed to the relative consistent audiovisual external stimuli received by participants during movie-watching. Higher BEN in the association cortex may be induced by the increased dynamic information processing and integration in the association cortex during movie-watching. Such processing is real-time and involves memory, emotion, and imagination which are subserved by medial-prefrontal cortex and parietal memory network (PMN) including lateral parietal cortex, and PCC [34]. Our functional decoding results using Neurosynth also indicate that the increased regional BEN induced by movie-watching is primarily associated with memory retrieval, autobiographical memory, and working memory. This suggests that during the processing of movie information, the integration of audiovisual information interferes with pre-existing internal mental states and memories and necessitates the engagement of working memory for information processing. The interference and modification of memory during movie-watching are also supported by research on other movie-watching. Baldassano et al revealed a nested hierarchy from short events in sensory regions to long events in AG and PCC and they proposed that brain activity is naturally structured into nested events, which form the basis of long-term memory representations [35]. Zadbood et al found that providing participants with information about the twist beforehand altered neural response patterns during movie-watching in the DMN and neural representations of past events encoded in the DMN are dynamically integrated with new information that reshapes our understanding in natural contexts [36]. The increase in BEN implies a reduction in long-range temporal coherence (LRTC) [18, 37], which is associated with long-term memory. He et al found that the variance and power-law exponent of the fMRI signal decrease during task activation, suggesting that the signal contains more long-range memory during rest and conversely is more efficient at online information processing during task [38]. During the resting-state, information is compressed through memory consolidation [39, 40], leading to lower BEN in DMN and PMN. However, long-term memory and internal mental state are disrupted and working memory is evoked during movie-watching. Our recent findings on regional BEN support this process. Our recent studies suggest lower resting regional BEN is associated with stronger task activation and deactivation in the cortex [26], while measurements of task-state BEN indicate higher BEN in the cortex relative to resting-state [41]. We also found consistently higher BEN in PCC during sad memory and ruminations compared to resting state across three different scanners [42]. The source of increased BEN may be related to break detailed balance during movie-watching, while the brain nearly obeys detailed balance when at rest [43]. Interestingly, movie-watching induced BEN patterns are supported by the previous inter-subject correlation (ISC) analysis [44], higher ISC in the sensory cortex, and lower ISC in the association cortex [45]. This is reasonable, as there is consistent audiovisual information input across participants, while individual internal mental states and memory are unique.

In conclusion, this study confirms the stability of regional BEN of brain activity induced by movie-watching. Furthermore, it establishes that BEN can effectively capture movie-related content, laying the groundwork for using BEN to characterize brain activity states across different movie stimuli and providing a new direction for exploring brain activity under natural paradigms using BEN in the future.

## Methods

### Dataset

#### Participants

All data were used in the study from the Human Connectome Project (HCP) 7T release. One hundred seventy-six (106 female) out of 184 subjects completed four rest state (REST) runs and four movie-watching (MOVIE) runs over two days, run1 (REST1, MOVIE1) and run2 (REST2, MOVIE2) in first day (Day1), run3 (REST3, MOVIE3), run4 (REST4, MOVIE4) in second day (Day2). All participants were healthy individuals between the ages of 22 and 36 years (mean age = 29.4, standard deviation = 3.3).

#### fMRI

All fMRI data were acquired on a 7 Tesla Siemens Magnetom scanner at the Center for Magnetic Resonance Research at the University of Minnesota. REST and MOVIE data were acquired using same gradient-echoplanar imaging (EPI) pulse sequence with the following parameters: repetition time (TR) = 1000 ms, echo time (TE) = 22.2 ms, flip angle = 45 deg, field of view (FOV) = 208 × 208 mm, matrix = 130 × 130, spatial resolution = 1.6 mm3, number of slices = 85, multiband factor = 5, image acceleration factor (iPAT) = 2, partial Fourier sampling = 7/8, echo spacing = 0.64 ms, bandwidth = 1924 Hz/Px. REST runs were always acquired first, followed by the movie runs in a fixed order, such that Day 1 consisted of REST1, REST2, MOVIE1, and MOVIE2, and Day2 consisted of REST3, REST4, MOVIE3, and MOVIE4, oblique axial acquisitions alternated between phase encoding in a posterior-to-anterior (PA) direction in REST1, REST3, MOVIE2 and MOVIE3, and an anterior-to-posterior (AP) phase encoding direction in REST2, REST4, MOVIE1 and MOVIE4.

During REST runs, participants were instructed to keep their eyes open with the relaxed fixation on a projected bright cross-hair on a dark background and presented in a darkened room. During MOVIE runs, participants passively viewed a series of video clips with audiovisual content. Each MOVIE run consisted of 4 or 5 clips, separated by 20 s of rest (indicated by the word “REST” in white text on a black background). Two of the runs, MOVIE1 and MOVIE3, contained clips from independent films (both fiction and documentary) made freely available under Creative Commons license on Vimeo. The other two runs, MOVIE2 and MOVIE4, contained clips from Hollywood films. The last clip was always a montage of brief (1.5 s) videos that were identical across each of the four runs to facilitate test-retest and validation analyses. For brief descriptions of each clip, see Table 1. The audio was delivered via Sensimetric earbuds. Each REST run was 900 TRs. MOVIE runs 1-4 were 921, 918, 915, and 901 TRs, respectively.

The fMRI images were preprocessed including motion correction, distortion correction, high-pass filtering, and nonlinear alignment to MNI template space plus regression of 24 framewise motion estimates and regression of confound time series identified via independent components analysis. Detailed information about data acquisition and preprocessing can be found in [46, 47]

### BEN mapping

The BEN mapping toolbox (BENtbx) [12] was used to calculate BEN using sam ple entropy (SampEn) [16]. The toolbox can be found at https://www.cfn.upenn.edu/zewang/BENtbx.php and https://github.com/zewangnew/BENtbx. More details of BEN calculation can be found in the original BENtbx paper [12] or our previous studies [18-20, 26, 48]. In this study, the window length was set to 3 and the cut-off threshold was set to 0.6 based on optimized parameters [12].

To avoid entropy calculation bias induced by the difference in the number of time points, and large changes in signal at the gap of individual clips, the first 10 time points were discarded for each clip [49]. Furthermore, we retained the longest three clips for each run, resulting in three movie clips per run, with each clip containing 170 TRs based on the shortest clip, and corresponding TRs were extracted from REST runs.

Similarly, we also computed the BEN maps from test-retest clips (83 TRs) at the end of each MOVIE run and corresponding TRs were extracted from REST runs to replicate and validate our analytical findings. We also included 900 TRs based on resting-state data length, followed by retaining the last 900 TRs during movie-watching for each run for calculating BEN. This was done to ascertain whether the BEN induced by movie-watching remains stable even in containing noise.

### Intraclass Correlation Coefficients

We assessed ICC between day 1, day 2 and all four resting state runs, and movie viewing runs separately. ICC was calculated using PyReliMRI [50] based on Shaefer 400 atlas [51].

### BEN differences between resting state and movie-watching

BEN maps of day 1, BEN maps of day 2 and four BEN maps of each subject were averaged to create the mean BEN maps for each subject, and paired sample T-tests were performed to determine the difference in BEN between movie-watching and the resting state.

### BEN differences between movie clips

One-way repeated measures analysis of variance (ANOVA) was employed to assess differences between the three movie clips or corresponding resting-state TRs within each run.

### Neurosynth Decoding

We employed Neurosynth (https://neurosynth.org) [52] to functionally decode the unthresholded statistical maps from the BEN difference between movie-watching and resting state. Detailed information about decoding can be found in [22].

## Acknowledgements

Data were provided by the Human Connectome Project, WU-Minn Consortium (Principal Investigators: David Van Essen and Kamil Ugurbil; 1U54MH091657) funded by th e 16 NIH Institutes and Centers that support the NIH Blueprint for Neuroscience Research; and by the McDonnell Center for Systems Neuroscience at Washington University.

## Data and code availability

All raw data are available at https://www.humanconnectome.org/hcp-protocols-ya-7t-imaging

BENtbx is available at https://www.cfn.upenn.edu/zewang/BENtbx.php.

Customed codes and further updates related to the study will be available at https://github.com/donghui1119/Brain_Entropy_Project/tree/main/Movie_BEN (upon publication o f the manuscript).

## CRediT authorship contribution statement

Dong-Hui Song: conceptualization, data analysis, visualization, manuscript drafting, and editing. Ze Wang: conceptualization, manuscript editing, supervision, project admi nistration.

